# The SapM phosphatase arrests phagosome maturation in an ESX-1 independent manner in *Mycobacterium tuberculosis* and BCG

**DOI:** 10.1101/2023.02.13.528419

**Authors:** Christian Xander, Saranathan Rajagopalan, William R Jacobs, Miriam Braunstein

## Abstract

*Mycobacterium tuberculosis* (*Mtb*) is an intracellular pathogen that survives and grows in macrophages. A mechanism used by *Mtb* to achieve intracellular survival is to secrete effector molecules that arrest the normal process of phagosome maturation. Through phagosome maturation arrest (PMA), *Mtb* remains in an early phagosome and avoids delivery to degradative phagolysosomes. One PMA effector of *Mtb* is the secreted SapM phosphatase. Because the host target of SapM, phosphatidylinositol-3-phosphate (PI_3_P), is located on the cytosolic face of the phagosome, SapM needs to be both released by the mycobacteria and escape the phagosome to carry out its function. To date, the only mechanism known for *Mtb* molecules to escape the phagosome is phagosome permeabilization by the ESX-1 secretion system. To understand this step of SapM function in PMA, we generated identical in-frame *sapM* mutants in both the attenuated *Mycobacterium bovis* bacille Calmette-Guérin (BCG) vaccine strain, which lacks the ESX-1 system, and *Mtb*. Characterization of these mutants demonstrated that SapM is required for PMA in both BCG and *Mtb*. Further, by establishing a role for SapM in PMA in BCG, and subsequently in a *Mtb* mutant lacking the ESX-1 system, we demonstrated that the role of SapM is ESX-1-independent. We further determined that ESX-2 or ESX-4 are also not required for SapM to function in PMA. These results indicate that SapM is a secreted effector of PMA in both BCG and *Mtb* and that it functions independent of the known mechanism for *Mtb* molecules to escape the phagosome.

## Introduction

Tuberculosis (TB) is a forgotten pandemic that continues to be a significant world health problem, with 1.6 million deaths from TB in 2021(1). The pathogenesis of *Mycobacterium tuberculosis* (*Mtb*), the bacteria responsible for TB, depends on the ability of *Mtb* to survive and replicate in macrophages(2, 3). To survive in macrophages, *Mtb* must subvert the antimicrobial immune responses of macrophages. One such way that *Mtb* promotes its survival in macrophages is by arresting the normal process of phagosome maturation (hereafter called phagosome maturation arrest (PMA)). As a result of PMA, *Mtb* avoids being delivered to acidic degradative phagolysosomes. Instead, *Mtb* remains in an early phagosome that is permissive for *Mtb* replication. Although it is central to *Mtb* pathogenesis, there remain significant gaps in knowledge of how *Mtb* achieves PMA. This is partly due to PMA being a complex process that relies on multiple *Mtb* effector molecules, many with redundant effects(2, 3). The redundancy highlights the importance of PMA to *Mtb* intracellular survival. It also means that elimination of a single effector may not necessarily eliminate PMA. All of the *Mtb* PMA effector proteins that are currently known are secreted from the bacteria as a necessary step for reaching their host targets. Consistent with this finding, the *Mtb* specialized SecA2 secretion system is required for PMA(4). Two SecA2-secreted proteins, SapM and PknG, contribute to PMA, but the data suggests the existence of at least one additional SecA2-secreted PMA effector(5).

SapM was the first secreted phosphatase discovered in *Mtb*(6). *In vitro* experiments with purified SapM, demonstrate SapM is capable of dephosphorylating phosphotidylinositols, with the highest specificity for phosphatidylinositol-3-phosphate (PI_3_P) and phosphatidylinositol-4,5-bisphosphate (PI(4,5)P_2_)(7, 8). PI_3_P is a regulatory lipid with a key role in phagosome maturation. During phagosome maturation, PI_3_P is generated on phagosomal membranes where it recruits PI_3_P-binding proteins, such as early endosome antigen 1 (EEA1) and the hepatocyte growth factor-regulated tyrosine kinase substrate (Hrs), which promote downstream fusion events between early phagosomes and late endosomes/lysosomes(9-13). Consistent with the ability of SapM to dephosphorylate PI_3_P and the role of PI_3_P in phagosome maturation, in *in vitro* assays purified SapM inhibits fusion between phagosomes and late endosomes(7). PI_3_P is located on the cytosolic-facing leaflet of the phagosome membrane(12, 14, 15). Therefore, for SapM to function in PMA the SapM protein must not only be secreted from *Mtb* but also escape the *Mtb-*containing phagosome in order to reach its PI_3_P target(16). In *Mtb*, the ESX-1 Type VII secretion system promotes phagosome permeabilization, which enable interactions between *Mtb* molecules and host cytosolic pathways(17-20). ESX-1 is also necessary for PMA(21). These properties have led to the idea that phagosome permeabilization by the ESX-1 system is the likely mechanism for effectors to reach the macrophage cytosol(2, 3). The mycobacterial lipid phthiocerol dimycocerosate (PDIM), which is also necessary for PMA, potentiates the membrane permeabilization effects of ESX-1(22-26).

The early studies of SapM did not involve *ΔsapM* mutants(6, 7); thus, a requirement for SapM during mycobacterial infection was not addressed until more recently. In *Mtb, ΔsapM* mutants are reported to be defective in PMA(27, 28). However, a *ΔsapM* mutant in the live attenuated TB vaccine *M. bovis* BCG strain, was reported to not exhibit a PMA defect(29). BCG is missing the region of difference 1 (RD1) of the *Mtb* genome, which encompasses genes encoding the ESX-1 system. Thus, the lack of a PMA defect for a BCG *ΔsapM* mutant was interpreted to reflect BCG SapM being unable to reach its cytosolic target, due to the absence of ESX-1 mediated phagosome permeabilization(2, 29). However, there are differences in the nature of the *Mtb* and BCG *ΔsapM* mutants studied and the methods used to monitor PMA(27, 28, 30). Therefore, to definitively address the issue of a required role of SapM in PMA as well as the role of the ESX-1 system in SapM function, we constructed identical in-frame *ΔsapM* mutants in *Mtb* and BCG and carried out side-by-side comparisons. Our results indicate that SapM is necessary for PMA in both *Mtb* and BCG and that SapM does not require the ESX-1 system to reach the macrophage cytosol to carry out its role in PMA. Our results also demonstrate that neither ESX-2 nor ESX-4 are required for PMA in the absence of ESX-1 (i.e., in BCG) arguing against the possibility that these other ESX systems provide the route for SapM to access the host cytoplasm.

## Materials and Methods

### Bacterial strains and media conditions

In this study, we used the *Mtb* H37Rv strain, and *ΔsecA2* and *ΔsapM* mutants generated in the H37Rv background (Suppl. Table 1). Additionally, we used the BCG Pasteur strain, and *ΔsecA2* and *ΔsapM* mutants generated in the BCG Pasteur background. *Mtb* and BCG strains were cultured using Middlebrook 7H9 (liquid) or 7H10 (agar) supplemented with 1x albumin dextrose saline (ADS), 0.5% glycerol, 0.05% Tween 80, and kanamycin (20μg/ml) or hygromycin (50μg/ml) when necessary(31).

### *ΔsapM* mutant construction in *Mtb* and BCG by allelic exchange using a counterselectable suicide plasmid

Unmarked *ΔsapM* mutants in *Mtb* H37Rv and BCG Pasteur were generated by two-step allelic exchange using a suicide counterselectable vector(31). The *ΔsapM* allelic exchange plasmid, pSEB1, was created using the hygromycin-resistant suicide *sacB* counterselectable plasmid pMP62(32). pMP62 was cut with SpeI and NheI and the resulting 8.8kb vector backbone fragment was used in a 3-part Gibson Assembly reaction with PCR products amplified from *M. tuberculosis* DNA upstream or downstream of *sapM*. The *sapM* upstream PCR product of 892 bp was amplified with PCR primer 5b [5’-gacgtatctagacacgtctgaaggctgaagtgctacttggagattc-3’] and primer 2 [5’-ttattggcggaccgcggagcatgccgggagtat-3’]. The *sapM* downstream PCR product of 751 bp was amplified with PCR primer 3 [5’-atgctccgcggtccgccaataaccgatatttggg-3’] and primer 4 [5’-ccatcccagctcggcaaggatcactagttgcccacctgcaaccagaagtc-3’]. The upstream and downstream PCR products were designed to have an overlapping region of sequence to promote 3-part Gibson assembly (NEB Gibson assembly kit) and primers 5b and 4 had extensions to anneal to vector pMP62. The resulting Δ*sapM* deletion is unmarked and encodes for a severely truncated SapM protein with the first 4 amino acids of SapM fused to the final 9 amino acids of SapM (MLRG-PPITDIWGD).

pSEB1 was electroporated into *Mtb* H37Rv or BCG Pasteur and the first recombination event was selected for on 7H10 plates supplemented with 50μg/ml hygromycin. Following growth in 7H9 media without antibiotics, bacteria that underwent a second recombination event between the deletion and wildtype alleles of *sapM* were selected by their growth on 7H10 plates supplemented with 3% sucrose to select for bacteria that lost the integrating *sacB* and hygromycin resistance containing plasmid. Sucrose-resistant and hygromycin-sensitive colonies were screened by PCR to identify *ΔsapM* deletion mutant versus wildtype recombinants. *ΔsapM* mutants were confirmed by immunoblot analysis of whole cell lysates with anti-SapM antibodies. Complementation strains were generated by electroporation of corresponding plasmids into mutant strains (Suppl. Table 2) as previously described(31).

### *ΔsapM/ΔeccD1 M. tuberculosis* double mutant construction by phage transduction

An ESX-1 (*ΔeccD1*) mutant was generated in the *ΔsapM Mtb* background to yield a *ΔsapM/ΔeccD1 Mtb* double mutant. The double mutant was constructed by specialized phage transduction using a temperature sensitive mycobacteriophage phAE159 to deliver a hygromycin marked *ΔeccD1* allelic exchange substrate (Suppl. Table 2), as described previously(4, 31). *Mtb* cultures were resuspended in MP phage buffer (50mM Tris/HCl pH7.5, 150mM NaCl, 10mM MgSO_4_, 2mM CaCl_2_) and high titer phage stock, prepared in *M. smegmatis*, added to a desired MOI of 10 phage to bacteria. The cell-phage mixture was incubated at 37-39°C for 4-6 hours and then plated onto 7H10 plates with hygromycin (50μg/ml) to select for allelic exchange mutants. Hygromycin-resistant colonies were screened by PCR to confirm mutants.

### *Δesx2 and Δesx4* BCG mutant construction by phage transduction

*esx-2* and *esx-4* operons, from r*v3884c* to r*v3895c* and *rv3342c* to *rv3452* respectively, were deleted in BCG Pasteur by specialized phage transduction (Suppl. Table 2), as described above. To make the *Δesx-2* and *Δesx-4* operon allelic exchange substrates, PCR reactions were performed using *Mtb* genomic DNA as a template. Forward and reverse flanking regions of *Mtb esx-2* and *esx-4* operons (1 kb each) were PCR amplified using primers ESX-2-1_LL [5’-ttttttttgcataaattgttccagccgcgtgggtcgc-3’], ESX-2-1_LR [5’- ttttttttgcatttcttgcaatgccatgaaatagctcgggatgtctcactgaggtctctagccgcatatcggctagtgcgg-3’], ESX-2-1_RL [5’- ttttttttgcatagattgcctcctaatgcgttaagctcacgagtgtctggtctcgtagtctcaataccccctggggctgc-3’], ESX-2-1_RR [5’-ttttttttgcatcttttgcagtatggtcgtggcgcagggg-3’], ESX-4_LL [5’- ttttttttccataaattggatcaatgcccgggcgataccca-3’], ESX-4_LR [5’- ttttttttccatttcttgggaaagtcttactgccgtgttgatgtctcactgaggtctctttccgggccccggagcgc-3’], ESX-4_RL [5’- ttttttttccatagattggaactcgtataacatccgcagcgagtgtctggtctcgtagctcccacggtggcgcgctg-3’], and ESX-4_RR [5’-ttttttttccatcttttggcggcaccggctcgccagc-3’]. Amplicons were either digested with Van91I or BstAPI based on the restriction site included in the primers and cloned into pYUB1471 backbone followed by ligation into phAE159 as previously described(33). The phages were propagated to high titers in *M. smegmatis* and verified by amplifying the flanking regions by PCR followed by Sanger sequencing. The *Δesx2 and Δesx4* BCG mutant strains were confirmed by PCR.

### Culture supernatant preparation

For preparing short term culture supernatants from *Mtb* and BCG, cultures were first grown to log-phase in supplemented 7H9 media. Cultures were then pelleted at 3,000rpm and washed once in PBS with 0.05% Tween 80 and then sub-cultured to a starting OD_600_ of 0.25 in fresh supplemented 7H9 media with 0.05% Tween 80 for 2 days. Cultures were pelleted at 3000rpm and supernatants were double filtered with 0.2μm filters to remove residual cells.

### Secreted SapM phosphatase activity assay

Secreted SapM phosphatase activity was measured as previously described(5, 34). Briefly, *Mtb* and BCG cultures were grown to mid-log phase and supernatants were normalized by culture OD_600_. Normalized filtered supernatants (160μl) were added in triplicate to a 96-well plate with 20μl 0.1mM Tris base pH 6.8 and 50mM p-nitrophenyl phosphate (pNPP) and 20μl 2mM sodium tartrate for a final volume of 200μl. SapM enzymatic activity is not inhibited by tartrate(6). Therefore, sodium tartrate was added to each sample to inhibit tartrate sensitive phosphatases. Samples were incubated at 37°C and the absorbance of the reactions were measured at OD_405_ every 3 minutes for 7 hours to calculate the rate of pNPP conversion, which was normalized to the amount of SapM secreted by wildtype *Mtb* or BCG.

### Macrophage infections

Bone-marrow derived macrophages (BMDMs) were isolated from femurs of C57/Bl6 mice by flushing with complete DMEM (DMEM supplemented with 10% fetal bovine serum, 5mM non-essential amino acids, and 5mM L-glutamine). Cells were then washed, resuspended, and plated in complete DMEM supplemented with 20% L-929 conditioned media (LCM) and incubated at 37°C with 5% CO_2_ for 6 days. Macrophages were then harvested using cold 5mM EDTA in PBS and seeded in complete DMEM supplemented with 10% LCM. For monitoring intracellular growth of *Mtb* or BCG, macrophages were seeded at 2×10^5^ macrophages/well in 8-well chambered slides. For LysoTracker microscopy experiments, macrophages were seeded at 1×10^5^ macrophages/well in 8-well chambered cover glass. After 24 hours, the macrophages were infected. For *Mtb*, macrophages were infected at MOI 0.1 for intracellular growth kinetics experiments or MOI 1 for LysoTracker staining experiments. For BCG, macrophages were infected at an MOI 1 for both growth kinetics experiments and LysoTracker staining. Prior to infection, the bacterial strains were grown to mid-log phase, washed twice using PBS containing 0.05% Tween 80, and then diluted in complete DMEM supplemented with 10% LCM. Macrophages were infected for 4 hours and then washed 3 times with PBS to remove extracellular bacteria. For growth kinetics experiments, cells were lysed using 0.1% Triton X-100 at various time points and lysates were diluted and plated on supplemented 7H10 agar for CFU enumeration.

### LysoTracker staining and co-localization

For LysoTracker staining, after 4 hours of infection followed by washing the infected macrophage monolayer to remove extracellular bacteria, DMEM complete media containing 100nM LysoTracker Red DND99 (Invitrogen) was added to each well and incubated at 37°C with 5% CO_2_ for 1 hour. The media was then removed and the chambered cover glasses were submerged and fixed in 4% paraformaldehyde (PFA) for 1 hour at room temperature. The fixed slides were then washed with PBS and Fluoromount-G (Southern Biotech) was added to each well to protect the fluorescent signal. Widefield microscopy was performed using an Olympus IX-81 and Metamorph software. Images were taken using a 40x oil objective. To visualize the *Mtb* or BCG, we used one of two methods. Either we took advantage of the endogenous autofluorescence of mycobacteria, using a CFP filter cube(4) or we stained the bacteria with DMN (4-*N,N*-dimethylamino-1,8-naphthalimide)-trehalose (DMN-Tre) obtained from OliLux Biosciences Inc.(35). For DMN-Tre staining, DMN-Tre was added to mycobacteria infected macrophages after the initial 4 hour infection and washes, stained for 24h, and subsequently stained with LysoTracker for 1 hour as previously described(36). A minimum of 10 images per well containing 250 mycobacteria-containing phagosomes were scored for LysoTracker co-localization. Two replicate wells (minimum of 500 phagosomes) were analyzed for each experiment and the data shown is representative of a minimum of two independent experiments.

### SapM antiserum production

To generate polyclonal antibodies against SapM, a peptide corresponding to amino acids 286-299 of SapM was synthesized. This peptide was injected into a specific-pathogen-free New Zealand white rabbits using TiterMax Gold adjuvant, at Pierce Custom Antibody Services (Rockton, IL). Antibodies were tested against wildtype *Mtb* and the *ΔsapM* mutant for specificity and purified by affinity chromatography.

### Immunoblots

10-25ml of bacterial cultures were pelleted and resuspended in extraction buffer (50mM Tris-HCl pH 7.5, 5mM EDTA, and 6% SDS) with protease inhibitor cocktail (Roche; 1mg/ml Aprotinin, 1mg/ml E-64, 1mg/ml Leupeptin, 50mg/ml Pefabloc SC, and 1mg/ml Pepstatin A). Bacterial whole cell lysates were made by bead beating twice for 90 seconds with <106μm glass beads using a MiniBeadbeater-16. Bacterial whole cell lysates, normalized by OD_600_ to load equal protein, were separated by SDS-PAGE and transferred to nitrocellulose membranes. Membranes were blocked and subsequently proteins were detected using anti-rabbit primary antibodies specific for SapM (1:10,000) or SatS (1:10,000)(34). An α-Rabbit IgG conjugated horseradish peroxidase secondary antibody (Bio-Rad) was used and the signal was detected using Western Lightning Plus-ECL chemiluminescent detection reagent (Perkin-Elmer). Alternatively, membranes were incubated with the α-Rabbit IRDye 800CW (LI-COR) and imaged using a LI-COR imager.

## Results

### *sapM* mutants of *Mtb* and BCG lack secreted SapM phosphatase activity

To study the role of SapM in *Mtb* PMA and intracellular growth in macrophages, we generated an in-frame, unmarked *ΔsapM* deletion mutant in the virulent H37Rv *Mtb* strain by two-step allelic exchange. In parallel, we generated an identical *ΔsapM* mutant in the *M. bovis* BCG Pasteur strain to enable comparison of the role of SapM in virulent *Mtb* and in the attenuated BCG vaccine. Successful deletion of *sapM* in both backgrounds was confirmed by PCR. Further validation of the *ΔsapM* mutants was carried out by immunoblot analysis on whole cell lysates using a rabbit polyclonal antibody raised against the last 14 amino acids of SapM (Fig. 1a), which demonstrated absence of SapM in the *ΔsapM* mutants compared to wildtype. In an operon and immediately downstream of *sapM* is the gene *satS*, which encodes a chaperone protein that stabilizes and promotes secretion of a subset of proteins by the SecA2 pathway, including SapM. SatS is also required for *Mtb* growth in macrophages(34). Thus, it was important to additionally confirm that the *ΔsapM* mutation did not disrupt *satS*. For this reason, whole cell lysates of *ΔsapM* mutants were additionally evaluated for SatS protein, using anti-SatS antibodies. Immunoblot analysis revealed SatS was present at wildtype levels in both *ΔsapM Mtb* and BCG mutants (Fig. 1a), confirming the *sapM* deletion did not disrupt downstream *satS* expression.

**Fig. 1.**
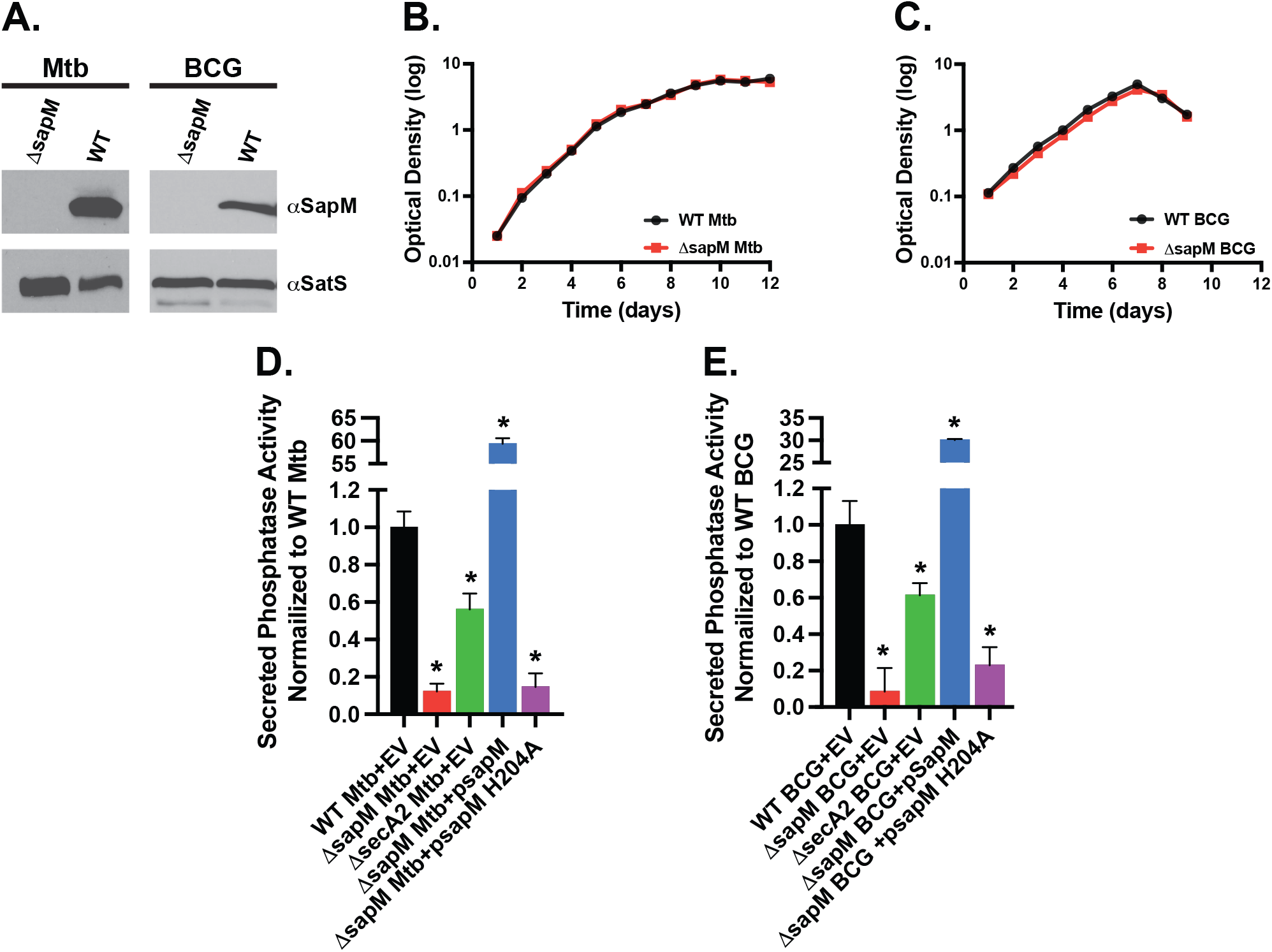
Δ*sapM* mutants in *Mtb* and BCG lack secreted phosphatase activity. (A) Equal protein from whole-cell lysates prepared from wildtype *Mtb* or BCG and corresponding Δ*sapM* mutants were examined for SapM or SatS protein by immunoblot with anti-SapM or anti-SatS antibodies. Δ*sapM* mutants in *Mtb* (B) or BCG (C) and their corresponding parental strains were grown in supplemented 7H9 media with 0.05% Tween 80 over 10-12 days. Optical density was used as a measure of in vitro growth. (D, E) Secreted phosphatase activity in culture supernatants from wiltype *Mtb* or BCG, corresponding Δ*sapM* or Δ*secA2* mutants, and complemented strains containing plasmids expressing wildtype SapM or a phosphatase-null mutant SapM_H204A_ were assessed using pNPP as a substrate, in triplicate. Activity was normalized to wildtype *Mtb* or BCG levels. * p<0.05 by ANOVA when compared to wildtype *Mtb* or BCG.

The *ΔsapM Mtb* and BCG mutants grew equally well as the wildtype parental strains *in vitro* in Middlebrook 7H9 media. Thus, at least under these conditions, SapM is not required for *in vitro* growth (Fig. 1b,c). We next examined filtered short-term culture supernatants prepared from wildtype, *ΔsapM* mutant, and complemented strains for secreted SapM-specific phosphatase activity using pNPP as a substrate. Because SapM phosphatase activity is resistant to sodium tartrate(6), sodium tartrate was included in the phosphatase assay to inhibit other secreted phosphatases and focus on SapM-specific activity. Both wildtype *Mtb* and BCG culture supernatants exhibited secreted phosphatase activity (Fig. 1d,e), while *ΔsapM Mtb* and BCG mutants exhibited a significant decrease in secreted phosphatase activity, as expected. The *ΔsapM* phosphatase defect could be complemented back by overexpressing wildtype *sapM* (*ΔsapM+psapM*) but not a phosphatase-null mutant allele *sapM*_*H204A*_(*ΔsapM+psapM*_*H204A*_)(5) in both *Mtb* and BCG *ΔsapM* mutants (Fig. 1d,e). In parallel, filtered culture supernatant from *ΔsecA2* mutants of *Mtb* and BCG, were also evaluated. As reported previously, SapM secretion was SecA2-dependent compared to wildtype. The observation that SapM phosphatase activity was only partially reduced in a *ΔsecA2* mutant, when compared to the defect observed in the *ΔsapM* mutant, is consistent with previous findings of some residual SapM secretion occurring in the absence of SecA2 through an unknown mechanism(5). These data demonstrated that the *Mtb* and BCG *ΔsapM* mutants constructed in this study lack secreted SapM tartrate-resistant phosphatase activity.

### SapM of *Mtb* is required for PMA and growth in a macrophage

Biochemical analysis of SapM demonstrates phosphatase activity on PI_3_P(4, 7, 8). Because PI_3_P drives phagosome maturation events, the PI_3_P dephosphorylation activity of SapM suggests a role of SapM in arresting phagosome maturation. Using our *ΔsapM* mutant of *Mtb* we directly addressed the requirement of SapM for PMA by *Mtb*. Murine bone marrow derived macrophages (BMDMs) were infected with wildtype, *ΔsapM* mutant, or complemented strains of *Mtb*. To assess PMA in infected macrophages, we used fluorescent LysoTracker, a dye that accumulates in acidified compartments, and fluorescence microscopy to determine *Mtb* localization to mature acidified phagosomes. As expected, the majority of wildtype *Mtb* did not localize to acidified (LysoTracker positive) phagosomes demonstrating the ability of *Mtb* to avoid/arrest the normal process of phagosome maturation. Approximately 20% of intracellular wildtype *Mtb* was observed in acidic phagosomes. In comparison to wildtype *Mtb*, the *ΔsapM Mtb* mutant exhibited a significantly higher association with acidified phagosomes indicating a requirement of SapM for *Mtb* PMA (Fig. 2a, 2b). The increase in localization to acidified phagosomes could be partially complemented by expressing wildtype *sapM* in the *ΔsapM Mtb* mutant (*ΔsapM+psapM*). When a phosphatase null *sapM*_*H204A*_ was expressed in the *ΔsapM Mtb* mutant (*ΔsapM+psapM*_*H204A*_) it did not complement the *ΔsapM Mtb* mutant defect in PMA, indicating the role of SapM in PMA specifically depends on its phosphatase activity. In parallel, we evaluated the maturation status of phagosomes containing the *ΔsecA2 Mtb* mutant. As reported previously, compared to wildtype the *ΔsecA2 Mtb* mutant exhibited a higher association with acidified phagosomes(4). Interestingly, the PMA defect of the *ΔsecA2 Mtb* mutant was significantly greater than that exhibited by the *ΔsapM Mtb* mutant. This observation is consistent with past studies indicating that SapM is not the only PMA effector secreted by the SecA2-pathway(5).

**Fig. 2.**
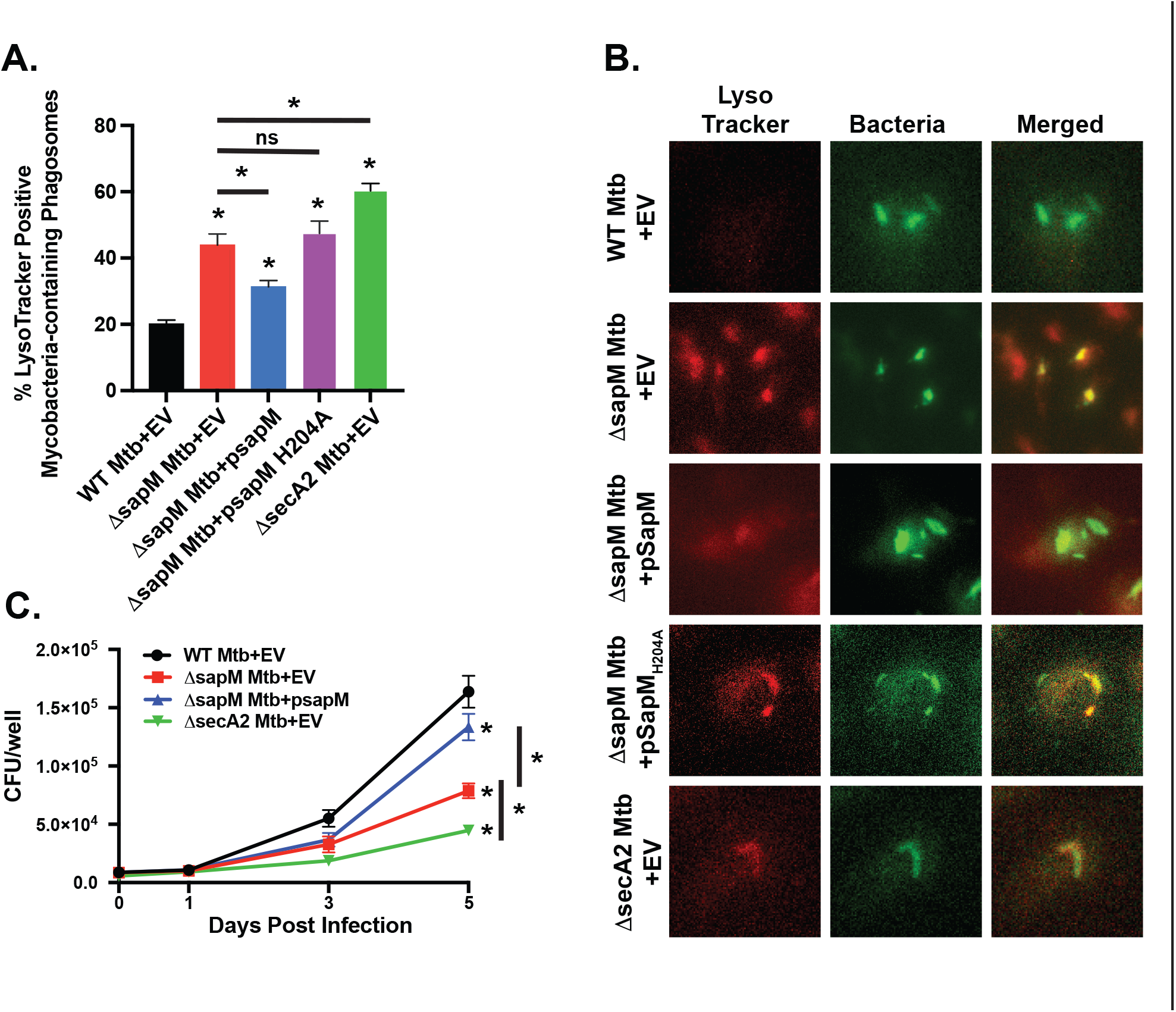
*Mtb* SapM arrests phagosome maturation and promotes growth in macrophages. BMDMs were infected with wiltype *Mtb*, Δ*sapM* or Δ*secA2* mutants containing empty vector plasmid (EV) or complemented strains containing plasmids expressing wildtype SapM or a phosphatase-null mutant SapM_H204A_. (A) The percentage of acidified mycobacteria-containing phagosomes at 1h post infection was determined by immunofluorescence microscopy using endogenous autofluorescence of mycobacteria and LysoTracker staining of quadruplicate wells. A minimum of 500 phagosomes were counted per condition. (B) Representative images used to quantify co-localization are shown. *Mtb* autofluorescence was re-colored green to highlight the co-localization in merged images. (C) BMDMs were infected at an MOI of 0.1 and intracellular growth was monitored over the course of 5 days. Triplicate wells were lysed and plated for CFU. Not significant (ns), * p<0.05 by ANOVA when compared to wildtype *Mtb* or between two strains connected by a line when otherwise noted.

As PMA is important for *Mtb* intracellular growth in a macrophage, we next determined if deletion of *sapM* in *Mtb* attenuates growth in a macrophage. BMDM were infected with the same strains as above and macrophage lysates were plated for colony forming units (CFU) over a five-day time course. The *ΔsapM Mtb* mutant was significantly attenuated for growth at five days post-infection when compared to wildtype and complemented strains (Fig. 2c) indicating a requirement for SapM in intracellular growth. Macrophages infected with the *Mtb ΔsecA2* mutant were evaluated in parallel. Consistent with the *ΔsecA2* mutant exhibiting a more severe defect in PMA and SapM not being the only SecA2-secreted PMA effector, the *ΔsecA2* mutant was significantly more attenuated than the *ΔsapM* mutant for intracellular growth in macrophages.

### SapM of BCG is required for PMA and growth in a macrophage

We next sought to determine if SapM is required for BCG to arrest phagosome maturation. As with the *Mtb* experiments, BMDM were infected with wildtype, *ΔsapM*, and complemented strains and PMA was determined by quantitating LysoTracker co-localization with BCG (Fig. 3a, Suppl. Fig. 1). In comparison to *Mtb*, BCG was not as efficient at PMA (wildtype BCG localized to more acidic compartments (approximately 40%) compared to wildtype *Mtb* (approximately 20%). This finding is consistent with previous reports that show BCG arrests phagosome maturation and with BCG being an attenuated strain of *M. bovis*(7, 16, 37). Nonetheless, even in the attenuated BCG strain background, a *ΔsapM* mutant exhibited a significantly higher association with acidic phagosomes compared to wildtype BCG (Fig. 3a, Suppl. Fig. 1). This defect in PMA arrest was partially complemented in the *ΔsapM+psapM* BCG complementation strain. This result indicates that SapM is also required for PMA in BCG. A *ΔsecA2* BCG mutant evaluated in parallel, also exhibited a defect in PMA. However, unlike with *Mtb*, BCG *ΔsapM* and *ΔsecA2* mutants showed equivalent defects in phagosome maturation arrest. Thus, in BCG it is possible that SapM is the only SecA2 secreted effector involved in arresting phagosome maturation.

**Fig. 3.**
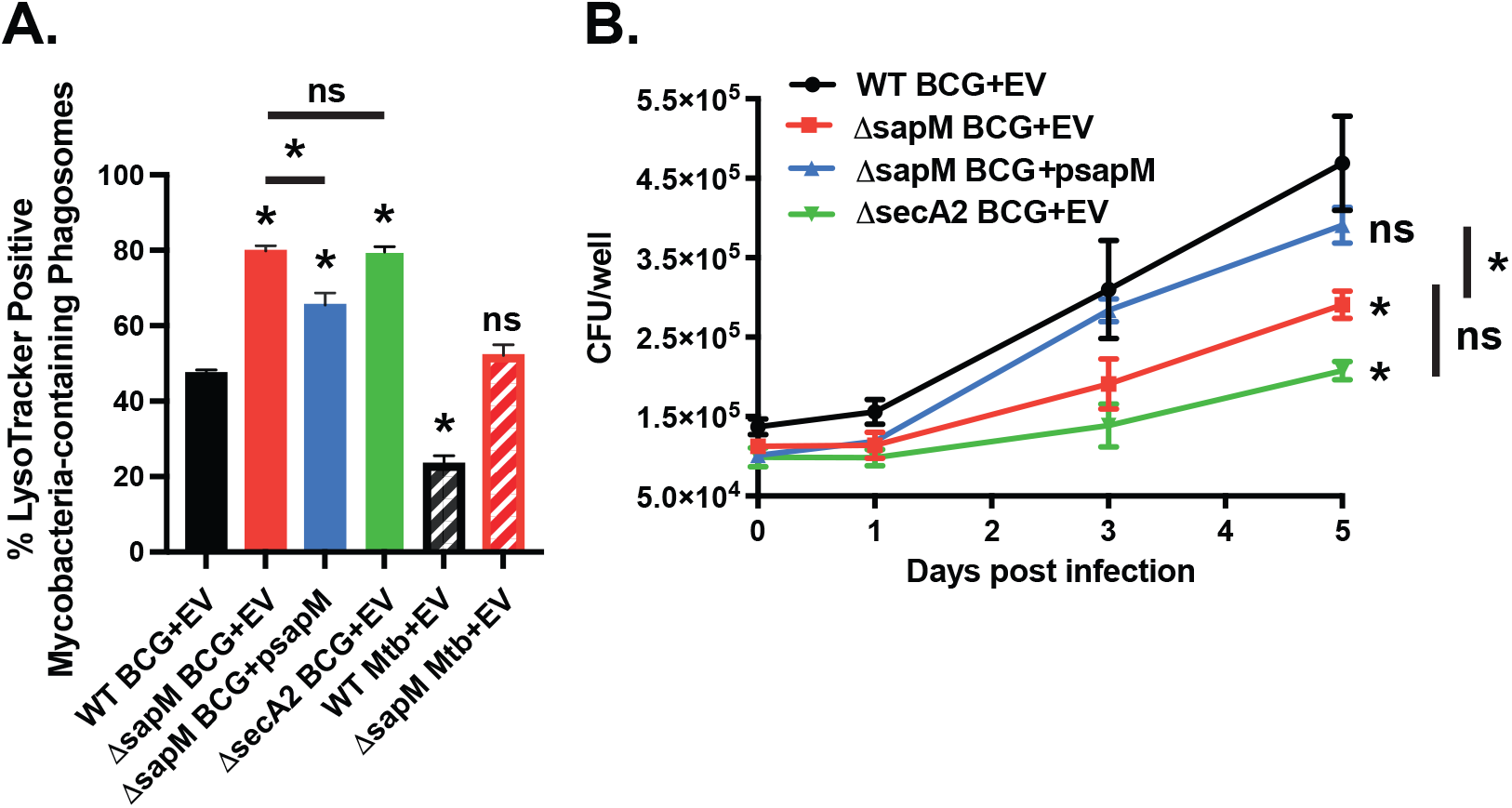
BCG SapM arrests phagosome maturation and promotes growth in macrophages. BMDMs were infected with wiltype BCG, Δ*sapM* or Δ*secA2* mutants containing empty vector plasmid (EV) or complemented strains containing a plasmid expressing wildtype SapM. (A) The percentage of acidified mycobacteria-containing phagosomes at 1h post infection was determined by immunofluorescence microscopy using endogenous autofluorescence of mycobacteria and LysoTracker staining of quadruplicate wells. A minimum of 500 phagosomes were counted per condition. (B) BMDMs were infected at an MOI of 1 and intracellular growth was monitored over the course of 5 days. Triplicate wells were lysed and plated for CFU. Not significant (ns), * p<0.05 by ANOVA when compared to wildtype BCG or between two strains connected by a line when otherwise noted.

Next, we determined whether SapM was required for BCG growth in BMDMs. Wildtype BCG grew in macrophages over five days albeit to a lesser degree than observed with *Mtb*. Compared to wildtype and the complemented *ΔsapM* BCG strain, the *ΔsapM* BCG mutant was significantly attenuated for intracellular growth (Fig. 3b). The *ΔsecA2* BCG mutant was also attenuated for growth in macrophages. However, as with the PMA defect, the *ΔsecA2* BCG mutant exhibited an intracellular growth defect that was not significantly different than that of the *ΔsapM* BCG mutant by ANOVA.

Taken together, the results from studying *ΔsapM Mtb* and BCG mutants are significant in demonstrating a conserved role for SapM in both *Mtb* and BCG in PMA and intracellular growth in a macrophage. However, a difference between *Mtb* and BCG may be the number of phagosome maturation arrest effectors and their dependence on the SecA2 pathway for secretion. In *Mtb* the SecA2 pathway appears to secrete multiple phagosome maturation arrest effectors that include SapM. However, the similarity in *ΔsapM* and *ΔsecA2* mutant phenotypes in BCG suggests that in BCG SapM may be the only SecA2 secreted effector acting to arrest phagosome maturation.

### The role of SapM in phagosome maturation arrest is ESX-1-independent

Because PI_3_P, the target of SapM and a factor promoting downstream phagosome maturation events, is located on the cytosolic leaflet of the phagosome(12, 14, 15), SapM must escape the phagosome to reach PI_3_P to play its role in PMA(16). The ESX-1 secretion system, which permeabilizes the phagosome membrane, is commonly assumed to be responsible for enabling secreted *Mtb* effectors to escape the *Mtb* containing phagosome and carry out their functions on cytoplasmic targets of the macrophage(2, 3, 17, 20). However, because BCG lacks RD1, which encompasses genes encoding the ESX-1 system, a role for SapM in BCG suggests that SapM reaches the macrophage cytoplasm in an ESX-1-independent manner (Fig. 3a). To determine if in *Mtb* the role of SapM is also ESX-1 independent, we deleted *eccD1* which encodes a critical component for a functioning ESX-1 system. We constructed an *ΔsapM/ΔeccD1 Mtb* double mutant and compared its ability to carry out PMA to that of wildtype *Mtb*, single *ΔsapM Mtb*, or single *ΔeccD1 Mtb* mutants in BMDM. As observed earlier, compared to wildtype *Mtb*, the single *ΔsapM Mtb* mutant exhibited a PMA defect (i.e., localized to significantly more acidic phagosomes) (Fig. 2a & 4a). The single *ΔeccD1 Mtb* mutant also exhibited a defect in PMA. However, the PMA defect **of** the single *ΔeccD1 Mtb* mutant was significantly greater than that of the single *ΔsapM Mtb* mutant, which is indicative of role(s) of ESX1 in PMA that are distinct from delivering SapM to the macrophage cytoplasm. Finally, the *ΔsapM/ΔeccD1 Mtb* double mutant exhibited the most severe PMA defect compared to either the single *ΔsapM* or *ΔeccD1 Mtb* mutants (Fig. 4a, Suppl. Fig. 2). The finding that the double mutant defect is greater than either single mutant demonstrates that SapM and ESX-1 have unrelated roles in PMA (i.e., the role of ESX-1 is not to deliver SapM to its cytoplasmic target). This result confirms that, as seen in BCG, SapM does not require ESX-1 to carry out its role in PMA. Introduction of wildtype *sapM* on p*sapM* to the *ΔsapM/ΔeccD1* mutant restored the *ΔsapM/ΔeccD1 Mtb* mutant phenotype back to the level of the single *ΔeccD1 Mtb* mutant, further demonstrating that SapM can perform its PMA function in the absence of ESX-1 (Fig. 4a, Suppl. Fig. 2). Together, our data in BCG and the *ΔsapM/ΔeccD1 Mtb* double mutant indicate the function of SapM in PMA is ESX-1-independent.

**Fig. 4.**
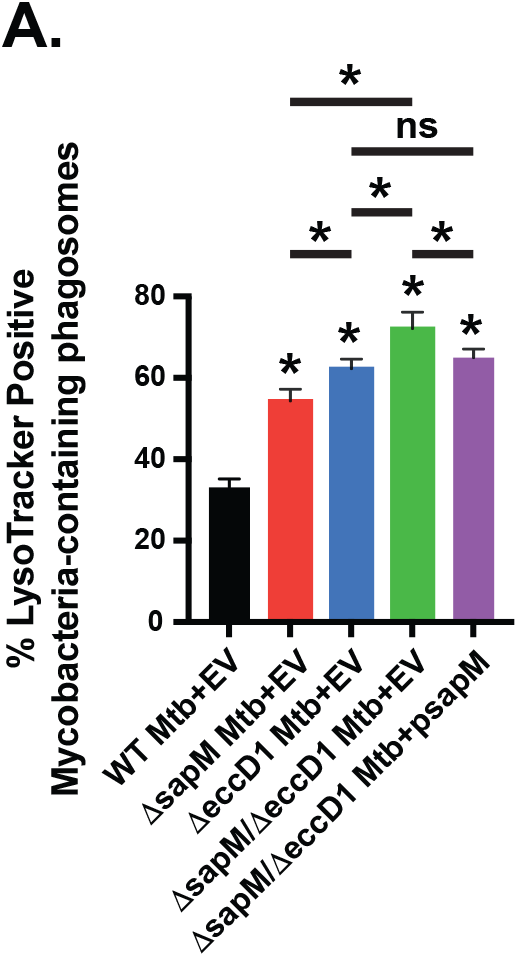
PMA by SapM is ESX-1-independent. BMDMs were infected with wiltype *Mtb*, Δ*sapM*, Δ*eccD1*, or Δ*sapM*/Δ*eccD1* mutants containing empty vector plasmid (EV) or complemented strains containing a plasmid expressing wildtype SapM. (A) The percentage of acidified mycobacteria-containing phagosomes at 24h post infection was determined by immunofluorescence microscopy using DMN-Tre and LysoTracker staining of quadruplicate wells. A minimum of 500 phagosomes were counted per condition. Not significant (ns), * p<0.05 by ANOVA when compared to wildtype *Mtb* or between two strains connected by a line when otherwise noted.

Previously, SapM was identified as one of the effector proteins secreted by the SecA2 pathway that account for the role of SecA2 in PMA(5). This was demonstrated using a *Mtb ΔsecA2* mutant strain that is engineered to have SapM secretion restored to equal or greater than wildtype levels. Restoration of SapM secretion in this strain is achieved by an ‘add-back’ strategy where SapM is overexpressed in the *ΔsecA2* mutant and the level of secreted SapM is restored, presumably by the unknown pathway responsible for residual SapM secretion seen in the absence of SecA2 (Fig. 1d, 1e). We additionally used the SapM ‘add-back’ strategy to confirm the ESX-1 independence of SapM function in PMA. As with the *ΔsecA2* mutant, when SapM was overexpressed and secretion restored to equal or greater than wildtype levels of SapM in the *ΔsecA2/ΔeccD1 Mtb* double mutant (*ΔsecA2/ΔeccD1+psapM*) a significant increase in PMA compared to the *ΔsecA2/ΔeccD1 Mtb* double mutant was observed (Suppl. Fig. 3). This data further strengthens our conclusion that the role of SapM in PMA is ESX-1-independent, and it implies the existence of a mechanism other than ESX-1 for SapM to escape the phagosome and reach its PI_3_P target in the macrophage.

### ESX-2 and ESX-4 are not required for the role of SapM in phagosome maturation arrest

A recent study suggests that ESX-2 and ESX-4 work in concert with ESX-1 to promote permeabilization of the *Mtb* containing phagosome membrane(38). Since SapM function in PMA was observed to be ESX-1-independent, we sought to determine if SapM cytosolic access, as a step in PMA, might be achieved by ESX-2 or ESX-4. To do this we generated *esx2* and *esx4* operon deletion mutants in BCG using a specialized phage transduction system. Successful mutants were confirmed by PCR. Because BCG lacks ESX-1, the resulting mutants were missing ESX-1 and ESX-2 or ESX-1 and ESX-4. We examined the ability of these mutants to block phagosome maturation compared to wildtype BCG and the *ΔsapM* BCG mutant with LysoTracker. As before, the *ΔsapM* BCG mutant localized to a significantly higher percentage of acidic phagosomes compared to wildtype BCG (Fig. 5a, 5b, Suppl. Fig. 4). However, the *Δesx2* BCG mutant showed no significant difference in PMA compared to wildtype BCG (Fig. 5a, Suppl. Fig. 4). Similarly, macrophages infected with the *Δesx4* BCG mutant did not reveal any significant difference in PMA compared to wildtype BCG (Fig. 5b). Together, these data suggest that neither ESX-2 or ESX-4 alone are responsible for the ESX-1 independent translocation of SapM to the cytosol for its role in PMA.

**Fig. 5.**
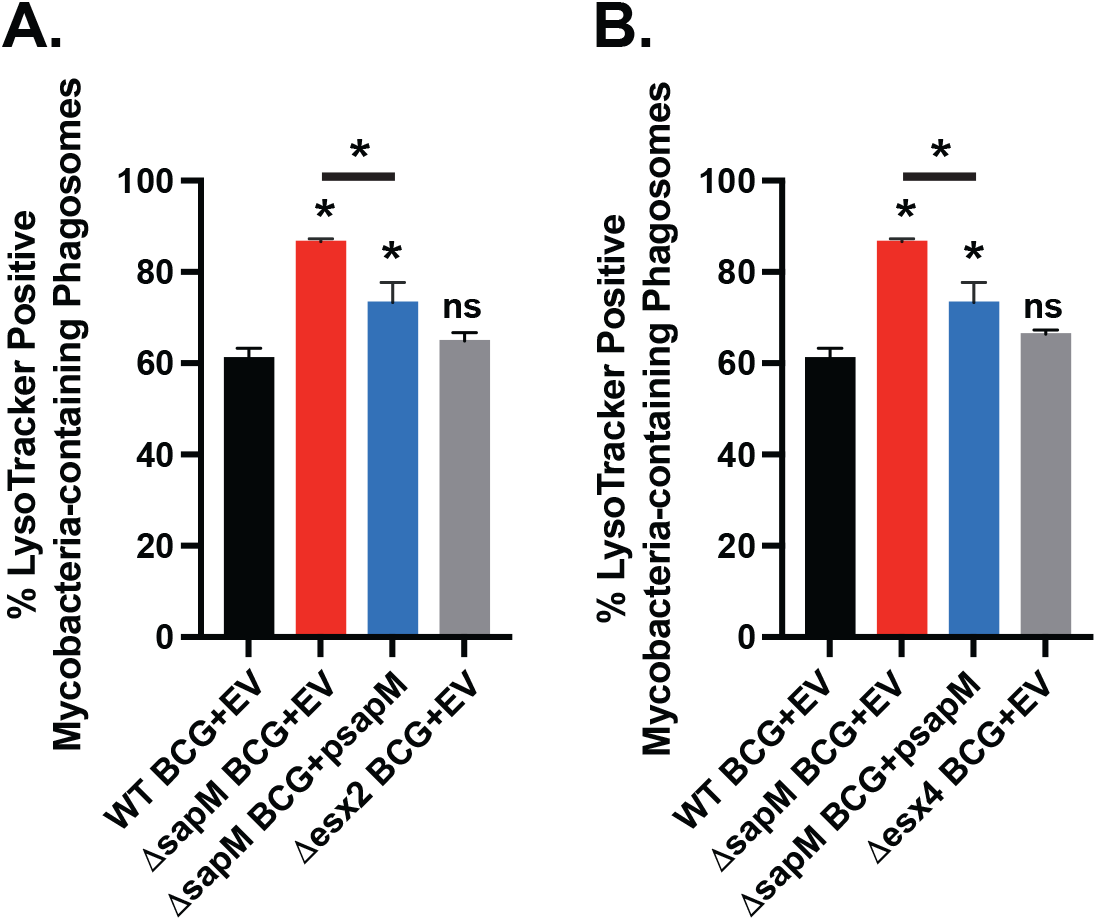
ESX-2 and ESX-4 are not required for SapM cytosolic access. BMDMs were infected with wiltype BCG, Δ*sapM*, Δ*esx2*, or Δ*esx4* mutants containing empty vector plasmid (EV) or complemented strains containing a plasmid expressing wildtype SapM. For ESX-2 (A) or ESX-4 (B), the percentage of acidified mycobacteria-containing phagosomes at 24h post infection was determined by immunofluorescence microscopy using DMN-Tre and LysoTracker staining of quadruplicate wells. A minimum of 500 phagosomes were counted per condition. Not significant (ns), * p<0.05 by ANOVA when compared to wildtype Mtb or between two strains connected by a line when otherwise noted.

## Discussion

Purified SapM is able to dephosphorylate phosphatidylinositol lipids *in vitro*, most notably phosphatidylinositol-3-phosphate (PI_3_P)(6-8). Normally, PI_3_P accumulates on phagosomes during phagosome maturation and phagosome-lysosome fusion(12). However, on *Mtb* and BCG-containing phagosomes PI_3_P levels are kept low (7). The discovery that SapM dephosphorylates PI_3_P *in vitro* suggested a role for SapM as an effector molecule of PMA. Later published studies of *sapM* mutants of *Mtb* supported a role for SapM in PMA. However, in BCG, which retains the ability to carry out PMA although at a reduced efficiency(4, 39, 40), evaluation of a *sapM* mutant did not lead to the same conclusion (i.e., a BCG *sapM* mutant was not defective in PMA)(29). BCG lacks the ESX-1 secretion system, which is required for PMA in *Mtb* and is often assumed to be the means by which *Mtb* effectors escape the phagosome. Therefore, the lack of an obvious role for SapM in BCG PMA was interpreted as reflecting an inability of BCG to transport SapM beyond the phagosome to act on PI_3_P(2, 27-29). However, in the past studies of *Mtb* and BCG *sapM* mutants, the nature of the mutants and methods for measuring PMA differed, which could have impacted the outcome of assessing the role of SapM. Because *sapM* is immediately upstream and in an operon with *satS* there was the potential for *sapM* mutations to have downstream effects on *satS*, which encodes a SecA2 chaperone that promotes secretion of SapM in addition to other SecA2-dependent proteins(34). For these reasons, here we sought to definitively determine the role of SapM in PMA in *Mtb* and BCG in a side-by-side comparison using identical in-frame, unmarked *sapM* mutations that do not impact *satS*.

Our results with an in-frame *Mtb ΔsapM* deletion mutant affirmed a role for SapM in *Mtb* in PMA and growth in macrophages (Fig. 2a, 2c). However, in contrast to published studies of a *sapM* transposon mutant in BCG(29), our evaluation of an in-frame *ΔsapM* mutation in BCG demonstrated a role for SapM in BCG PMA and growth in macrophages (Fig. 3a, 3b). Given this result, it is noteworthy that in the early report by Vergne *et al*.(7) they observed infection with live BCG to be associated with PI_3_P removal from the phagosome membrane in a manner consistent with the activity of SapM. Importantly, the PMA and intracellular growth defects of the *ΔsapM* mutants could be complemented by introduction of plasmids expressing wildtype *sapM* but not by a catalytically inactive *sapM*_*H204A*_ allele (Fig. 2a). These results demonstrate that SapM has a conserved role in PMA and intracellular growth in *Mtb* and BCG.

The kinase that converts phosphatidylinositol to PI3P and PI3P itself is located on the cytosolic-facing leaflet of the phagosome membrane(12, 14). Therefore, for SapM to dephosphorylate PI_3_P it is critical that SapM reach the cytosolic side of the phagosomal membrane. Such a requirement raises the question of how secreted SapM reaches the macrophage cytosol. The ESX-1 secretion system of *Mtb* is required for PMA(4, 21, 41) and, working with PDIM, it is known to permeabilize the phagosome membrane to allow interactions between *Mtb* products and host cytosolic molecules, including innate immune sensors(18, 19). Thus, it is generally assumed that secreted *Mtb* effector proteins will exit the phagosome with the assistance of the ESX-1 secretion system(2, 3, 17, 20). However, currently there are only two *Mtb* proteins, Tuberculosis Necrotizing Toxin (TNT) and Mpt64, demonstrated to require ESX-1 for their delivery to the macrophage cytosol and neither protein has a known role in PMA(38, 42). In the study presented here, we provide multiple lines of evidence to indicate that SapM does not depend on ESX-1 to reach its host cell target and function in PMA. First, we demonstrated a role for SapM of BCG in PMA even though BCG lacks a functional ESX-1 system (Fig. 3a). Second, with a *sapM* complementation plasmid and an *ΔsapM/ΔeccD1* double mutant in *Mtb* we demonstrated SapM to function in PMA in the absence of ESX-1 in *Mtb* (Fig. 4a). Further, the *ΔsapM/ΔeccD1* double mutant exhibited a more pronounced PMA defect than either an *ΔeccD1* or *ΔsapM* single mutant (Fig. 4a). The more severe phenotype of the *ΔsapM/ΔeccD1* double mutant indicates that ESX-1 and SapM have independent functions in PMA, and it argues against ESX-1 delivering SapM to its host cell target for its role in PMA. Third, in our ‘add-back’ system, which was used previously to study the contribution of secreted SapM to the role of the SecA2 pathway in macrophages, we also observed SapM to impact PMA in an ESX-1 independent manner (Suppl. Fig. 4). Together, these results provide strong evidence for SapM reaching its host target in PMA in an ESX-1 independent manner. This conclusion is also supported by a study performed with *Mycobacterium marinum* in which phosphatase overexpression (SapM, PtpA, and PtpB) is associated with reduced phagosome PI_3_P levels in an *esx-1* mutant background (43). Thus, the idea that all mycobacterial effectors rely on ESX-1 to confer cytosolic access and activity on host targets needs to be reconsidered. Further, the role for ESX-1 in PMA remains to be understood as PMA effectors that do depend on ESX-1 for their activity remain to be identified (3). Interestingly, partial inhibition of phagosome acidification (*i*.*e*., maturation) is proposed to be a prerequisite for ESX-1 mediated phagosome permeabilization(44). This raises the interesting possibility that ESX-1 independent transport of SapM could promote initial reduction of phagosome acidification as a step towards ESX-1 permeabilization and robust PMA.

Recently, *ΔeccC2* or *ΔeccC4* mutants of *Mtb*, which encode ATPases essential for the ESX-2 and ESX-4 secretion systems, were shown to have phagosome permeabilization defects(38). Thus, suggesting that ESX-2 and ESX-4 systems work with ESX-1 to permeabilize the phagosome. Therefore, we explored the possibility that ESX-2 or ESX-4 secretion systems account for SapM escape from the phagosome. To address this possibility, we constructed deletion mutants in which the entire *esx2* or *esx4* operon sequences were deleted in BCG. Due to the absence of ESX-1 in BCG the resulting mutants can be considered *Δesx1/Δesx2* or *Δesx1/Δesx4* double mutants. The level of PMA observed in macrophages infected with these strains was no different than that associated with the parental BCG strain. Thus, BCG does not require ESX-2 or ESX-4 for PMA thereby ruling out the possibility that these other ESX systems are responsible for the ESX-1 independent SapM delivery to its host cell target in PMA. The question of how SapM gains cytosolic access to dephosphorylate PI_3_P remains to be answered. SapM belongs to the phospholipase C/phosphatase superfamily which includes the *Francisella tularensis* acid phosphatase AcpA(8, 45). AcpA is involved in disrupting the phagosome membrane to allow *F. tularensis* cytosolic access(46). Thus, it is possible that the SapM protein may enable its own escape from the phagosome to allow it to act on phagosomal PI_3_P. Alternatively, SapM may gain cytosolic access via an unknown mechanism that we have yet to discover.

Since we previously identified SapM as one of the *Mtb* effectors secreted by the SecA2 pathway, we also compared the phenotypes of *ΔsapM* and *ΔsecA2* mutants in our studies. In both PMA and intracellular growth, the *Mtb ΔsecA2* mutant exhibited a more severe phenotype than the *ΔsapM* mutant. This finding is consistent with our past studies that indicated that the *Mtb* SecA2 pathway secretes multiple PMA effectors (5). One of these other effectors is PknG but additional effectors also exist. However, unlike in *Mtb*, in BCG there was no significant difference in PMA or intracellular growth phenotypes between *ΔsecA2* and *ΔsapM* mutants. One possible explanation is that in BCG SapM is the only SecA2-dependent effector involved in PMA.

In conclusion, here we demonstrated that SapM is required for PMA and growth in macrophages in both *Mtb* and BCG. It is noteworthy that *ΔsapM* mutants of BCG and *Mtb* are being studied for their potential to be used as live-attenuated TB vaccines(27, 29, 30, 47). Thus, the knowledge that a role for SapM in PMA is conserved in both species may help explain how *ΔsapM* mutations improve antigen presentation and vaccine efficacy. We also demonstrated that the function of SapM in PMA does not depend on ESX-1, nor does it depend on ESX-2 or ESX-4. Thus, our data reports on the existence of a mechanism other than ESX-1 mediated phagosome permeabilization as a way for *Mtb* effectors to escape the phagosome and reach host targets.

## Acknowledgements

We gratefully acknowledge Sebastian Murcia for assistance in constructing the *ΔsapM* allelic exchange plasmid, Kate Zulauf for constructing the *ΔsecA2/ΔeccD1* and complemented strains, and the Microscopy Services Laboratory at UNC for microscopy advice and use of their microscopes. MB acknowledges support of AI149727 and WRJ acknowledges support of AI026170 and AI156853.

## Supplemental Figure Legends

**Suppl. Fig. 1. Representative microscopy images for *ΔsapM* BCG LysoTracker experiments**. BCG infected BMDMs where stained with LysoTracker for 1 hour. Representative images used to quantify co-localization in Fig. 3 are shown. BCG autofluorescence was re-colored green to highlight the co-localization in merged images.

**Suppl. Fig. 2. Representative microscopy images for *ΔsapM*/*ΔeccD1 Mtb* LysoTracker experiments**. *Mtb* infected BMDMs where stained with LysoTracker for 1 hour. Representative images used to quantify co-localization in Fig. 4 are shown. *Mtb* were stained with DMN-Tre to highlight the co-localization in merged images.

**Suppl. Fig. 3. SapM “add back” experiment demonstrates ESX-1-independent function of SapM**. (A) Secreted phosphatase activity in culture supernatants from wildtype *Mtb*, corresponding *ΔsecA2* or *ΔsecA2/ΔeccD1* mutants, and ‘add back’ strains expressing wildtype SapM were assessed using pNPP as a substrate, in triplicate. BMDMs were infected with wiltype *Mtb, ΔsecA2*, or *ΔsecA2/ΔeccD1* mutants containing empty vector plasmid (EV) or strains overexpressing wildtype SapM on a plasmid. (B) The percentage of acidified mycobacteria-containing phagosomes at 24h post infection was determined by immunofluorescence microscopy using DMN-Tre and LysoTracker staining of quadruplicate wells. A minimum of 500 phagosomes were counted per condition. Not significant (ns), * p<0.05 by ANOVA when compared to wildtype *Mtb* or between two strains connected by a line when otherwise noted. (C) Representative images used to quantify co-localization are shown. *Mtb* were stained with DMN-Tre to highlight the co-localization in merged images.

**Suppl. Fig. 4. Representative microscopy images for *Δesx2* and *Δesx4* BCG LysoTracker experiments**. (A) *Δesx2* or (B) *Δesx4* BCG infected BMDMs where stained with LysoTracker for 1 hour. Representative images used to quantify co-localization in Fig. 5 are shown. BCG were stained with DMN-Tre to highlight the co-localization in merged images.

